# Figra: A WebAssembly-based Excel Add-in for publication-quality scientific visualization with ggplot2

**DOI:** 10.64898/2026.05.06.723320

**Authors:** Yoshiaki Sato

**Affiliations:** Department of Medicine, Keck School of Medicine of University of Southern California (USC), Los Angeles, CA 90033, USA; Norris Comprehensive Cancer Center, Keck School of Medicine of USC, Los Angeles, CA 90033, USA

## Abstract

Data visualization is a critical step in scientific communication. Most researchers rely on subscription-based software for this purpose, which requires ongoing licensing costs. Free alternatives such as R and Python offer publication-quality output but demand programming expertise that many researchers do not possess. Artificial intelligence tools can assist with figure generation but remain frustrating when users wish to fine-tune specific visual parameters to their preference. Meanwhile, Microsoft Excel, the most widely used tool for scientific data storage and management, offers limited visualization capabilities, forcing researchers to transfer their data to external software as an extra step before creating figures. Here we present Figra, a free Excel Office Add-in that eliminates this extra step by enabling publication-quality ggplot2-based figure generation directly within Excel, with simple and direct control over every visual option. Figra leverages WebAssembly technology (webR) to execute R code entirely within the browser, requiring no R installation, no subscription, and no server connection. The add-in supports over 20 chart types spanning distribution plots, grouped comparisons, time-series, scatter plots, and specialized curve-fitting analyses. For applicable chart types, Figra performs automated or manual statistical analysis supporting both paired and unpaired designs across two or more groups. Additionally, Figra exports simplified, executable R code that reproduces the displayed figure, serving as an educational tool for researchers wishing to learn ggplot2. Figra is open-source and freely available at https://h20gg702.github.io/figra-pages/index.html while the source code is provided at https://github.com/h20gg702/Figra.

## Introduction

Data visualization is a crucial component of the scientific workflow, facilitating data exploration, extraction of experimental findings, and communication of results^1,2^. Most life scientists collect and store their experimental data in Microsoft Excel, yet Excel’s native charting capabilities are insufficient for publication-quality figures, forcing researchers to transfer their data to a separate tool before they can create a single graph. This extra step is not only inefficient but leaves the visualization workflow largely undocumented and difficult to reproduce.

The most common solution is subscription-based software such as GraphPad Prism, which offers an intuitive interface but requires ongoing licensing costs. Free alternatives such as R with ggplot2 produce exceptional figures and provide full reproducibility, but demand coding expertise that many wet laboratory researchers do not possess. Web-based tools such as BoxPlotR^3^ and PlotsOfData^4^ have lowered this barrier but offer limited chart types and customization, and require data upload to external servers. More recently, artificial intelligence tools can assist with figure generation, but iteratively refining visual parameters through prompting remains frustrating compared to direct graphical interaction.

Here we present Figra, a free Excel Office Add-in that eliminates the gap between data storage and figure creation. Users load their Excel data, configure their figure through a simple graphical interface, and insert publication-quality ggplot2-based figures directly back into their worksheet, with no coding, no subscription, and no data upload required. Figra leverages webR^5^ to execute R code entirely within the browser, supports 24 chart types with integrated statistical analysis, and exports executable R code for each figure as a reproducible record of the visualization workflow.

## Results

Figra is implemented as a Microsoft Excel Office Add-in presenting a Figra control panel organized into nine tabs: Data & Chart, Converter, Colors & Style, Text & Font, Theme & Layout, Statistics, Analysis, and Help. Users select their data range in Excel, load it into Figra, configure the figure through graphical controls, preview the result, and insert the finished figure directly back into their worksheet, the entire workflow takes place without leaving Excel.

Figra supports 24 chart types spanning single-group and grouped comparisons across a wide range of visualization styles (Figure 1, Table 1). Single-group charts include histogram, box plot, violin plot, dot plot, scatter plot, bar chart, and line chart. Grouped variants allow multi-group comparison for box, violin, bar, and line charts. All chart types support extensive visual customization including five font families, eight ggplot2 themes, linear and logarithmic axis scales, custom colors with adjustable transparency, axis label rotation, and flexible export dimensions and resolution, all accessible through direct graphical controls without any coding. Visual style settings can be saved as named presets and reloaded across sessions, facilitating consistent figure styling throughout a manuscript (Supplementary Table S1).

**Table 1.** Chart types supported by Figra and availability of integrated statistical analysis.

| Chart type | Single | Grouped | Dot overlay | Pre-calc error bars | Statistical analysis |
| --- | --- | --- | --- | --- | --- |
| Histogram | ✓ | — | — | — | — |
| Box plot | ✓ | ✓ | ✓ | — | ✓ |
| Violin plot | ✓ | ✓ | ✓ | — | ✓ |
| Dot plot | ✓ | — | — | — | — |
| Scatter plot | ✓ | — | — | — | — |
| Bar chart | ✓ | ✓ | ✓ | ✓ | ✓ |
| Line chart | ✓ | ✓ | , | ✓ | ✓ |
| IC50 dose-response | ✓ | ✓ | — | — | — |
| LQ survival curve | — | ✓ | — | — | ✓ |

**Table 2.** Example input data formats accepted by Figra. Figra accepts two input formats depending on the chart type selected.

| Category | Values |
| --- | --- |
| shControl | 1.00 |
| shControl | 1.15 |
| shControl | 0.98 |
| shGeneA-a | 0.61 |
| shGeneA-a | 0.55 |
| shGeneA-a | 0.49 |
| shGeneA-b | 0.54 |
| shGeneA-b | 0.60 |
| shGeneA-b | 0.57 |

| Category | Mean | Error |
| --- | --- | --- |
| shControl | 1.04 | 0.09 |
| shGeneA-a | 0.55 | 0.06 |
| shGeneA-b | 0.57 | 0.03 |

For grouped chart types, a Group column is added as the first column (e.g., Group | Category | Values for grouped bar). For line and dose-response charts, the Category column is replaced by a continuous X-axis variable (e.g., time or concentration). The built-in wide-to-long data converter (Converter tab) can reshape data from wide format (one column per group) to the long format required by Figma.

**Figure 1.**
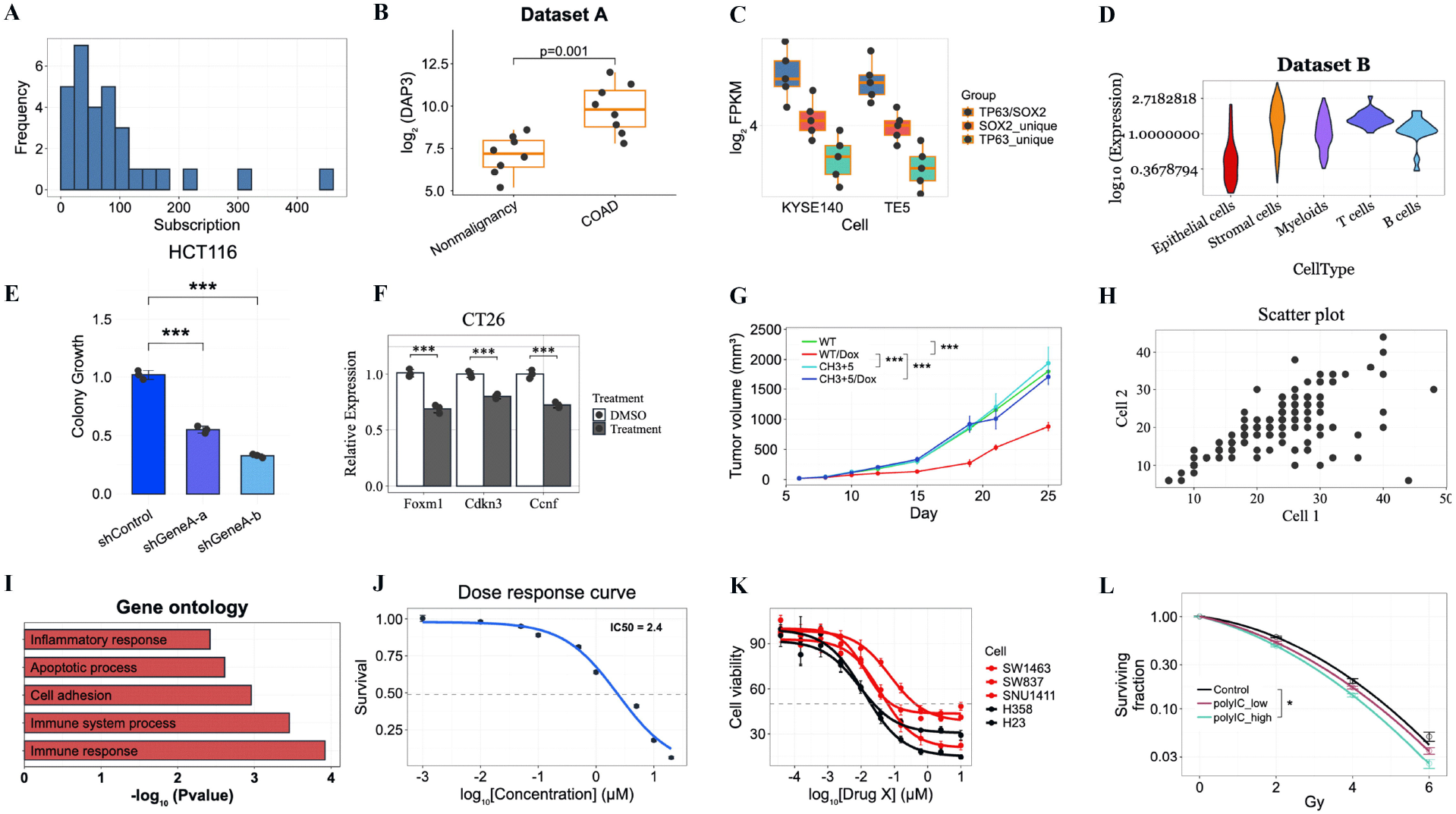
Capability showcase of Figra. (A) Histogram of a single continuous variable. (B) Box plot with superimposed data points comparing two groups; Figra automatically selected the appropriate statistical test and annotated significance (p = 0.001). (C) Grouped box plot with dots for two cell lines across three conditions. (D) Violin plot with per-category coloring and a log_10_-transformed Y axis. (E) Bar chart with error bars and individual data points for three groups (HCT116); Dunnett’s post-hoc test versus the control group with p-values annotated directly on the figure. (F) Grouped bar chart with error bars comparing two conditions across three targets. (G) Grouped line chart with error bands for four groups over time; pairwise statistical comparisons are displayed with bracket annotations. (H) Scatter plot for two continuous variables. (I) Horizontal bar chart for ranked categorical data. (J) Single-group dose–response curve fitted with a 4PL model; IC_50_ is automatically calculated and annotated (IC_50_ = 2.4 μM). (K) Grouped dose–response curves for multiple conditions fitted simultaneously with the 4PL model. (L) Linear-quadratic clonogenic survival curves for three groups; α, β, α/β ratio, SF_2_, D_10_, and D_0_ are automatically calculated and exported.

For chart types that include raw data points, Figra performs fully automated statistical analysis with appropriate test selection based on data characteristics or manual statistical analysis based on user’s knowledge (Figure 2, Table 1). The pipeline begins with normality assessment using the Shapiro-Wilk test. For unpaired designs, normally distributed data proceed to variance testing followed by Student’s t-test or Welch’s t-test for two groups, or ANOVA for three or more groups. Non-normally distributed data are analyzed using the Wilcoxon rank-sum test or Kruskal-Wallis test. For paired designs, Figra applies the paired t-test or repeated measures ANOVA for normally distributed differences, and the Wilcoxon signed-rank test or Friedman test otherwise. Post-hoc testing supports five methods: Tukey HSD, Bonferroni, Holm, Dunnett, and Dunn. Statistical significance is annotated directly on the figure using stars, compact letter display, or exact p-values according to user preference.

**Figure 2.**
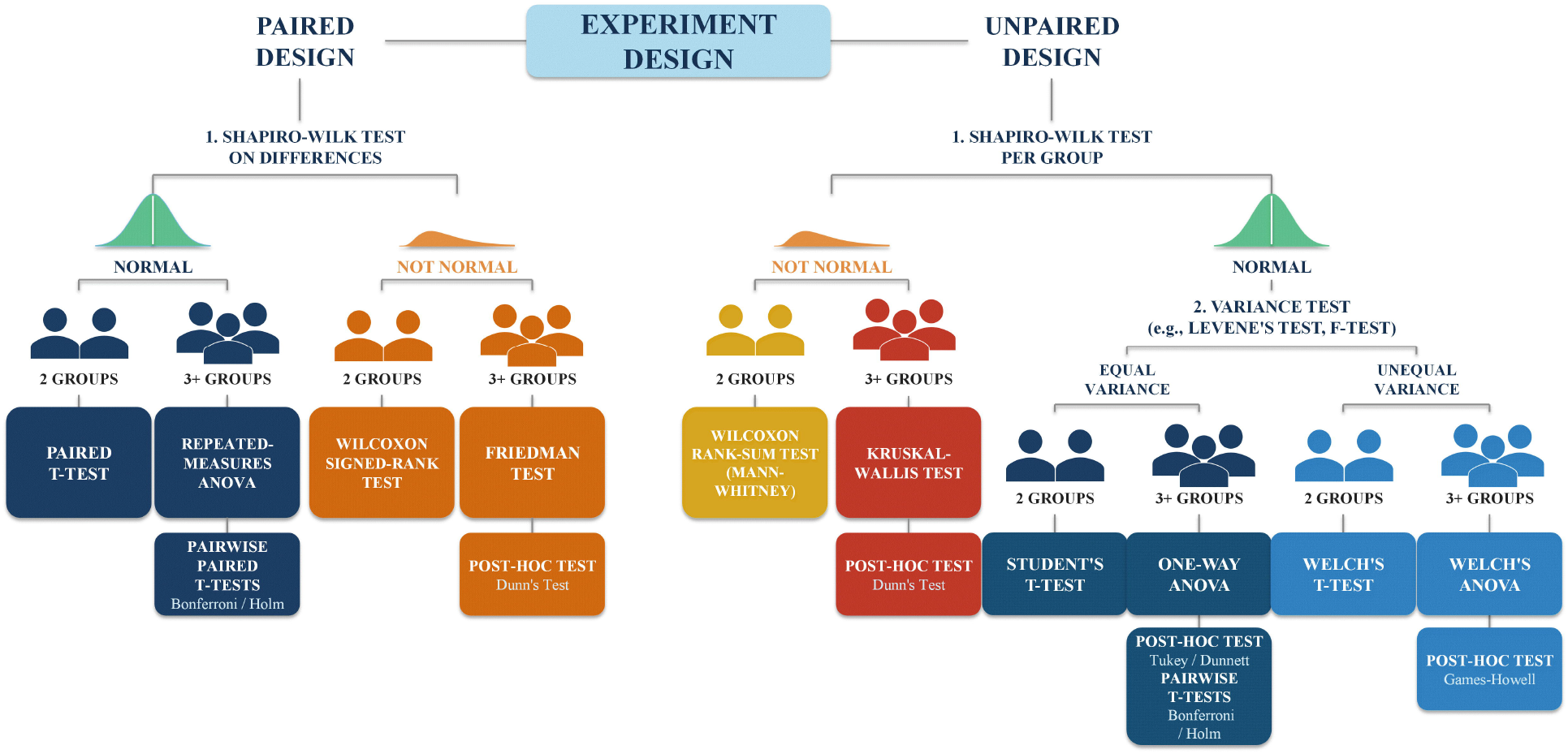
Automated statistical test selection in Figra. The flowchart illustrates the decision logic applied when the “Show statistical significance” option is enabled. For each dataset, Figra first determines whether measurements are paired or unpaired. For paired data, normality is assessed on the pairwise differences using the Shapiro–Wilk test; the appropriate paired test (paired t-test or Wilcoxon signed-rank) or repeated-measures procedure (repeated-measures ANOVA or Friedman test) is then selected automatically. For unpaired data, normality is assessed per group; if normality holds, a variance test (Levene or F-test) further guides selection between equal-variance and unequal-variance procedures (Student’s t-test / one-way ANOVA vs. Welch’s t-test / Welch’s ANOVA). Non-normal data are analyzed with distribution-free tests (Wilcoxon rank-sum or Kruskal–Wallis). For experiments with three or more groups, post-hoc testing is performed using user-selected methods: Tukey HSD, Games-Howell, Bonferroni correction, Holm correction, Dunnett’s test, and Dunn’s test are supported. Statistical results are rendered as significance symbols (stars), compact letter displays, or exact p-values according to user preference.

Figra also supports two specialized curve-fitting analyses (Figure 1I–K). For dose-response data, Figra fits the four-parameter logistic model, 4PL model, (Response = Bottom + (Top − Bottom) / (1 + (IC_50_/Dose)^Hill)) to the data and reports IC50, Hill coefficient, and upper and lower asymptotes (Top, Bottom) per group, with support for simultaneous fitting across multiple conditions^6^. For radiation clonogenic survival data, Figra fits the linear-quadratic model (SF = exp(−*α·*D − β·D^2^)) to the data and reports α, β, α/β ratio, SF2 (surviving fraction at 2 Gy), D_10_ (dose for 10% survival), and D_0_ (dose for 37% survival) per group^7^.

To accept a variety preference of data saving format, Figra includes a built-in wide-to-long data format converter, allowing users to reshape their spreadsheet data directly within Excel without external tools (Supplementary Table S2). Additionally, for each figure generated, Figra exports simplified executable R code that reproduces the displayed figure, serving both as a reproducible record of the visualization workflow and as an educational resource for researchers wishing to learn R.

For chart types that perform statistical analysis, Figra can export the full statistical results, including summary statistics, normality test results, test selection rationale, and post-hoc comparisons, directly into selected Excel cells alongside the data. For specialized curve-fitting analyses, Figra similarly exports model parameters and fitted curve data. Representative examples of this output are provided in Supplementary Tables S3–S6.

## Conclusion

Figra democratizes publication-quality scientific visualization by bringing the ggplot2 ecosystem directly into Microsoft Excel through WebAssembly technology, eliminating the extra step between data storage and figure creation. With support for 24 chart types, comprehensive customization, automated statistical analysis for paired and unpaired designs, specialized curve-fitting tools, and educational R code export, Figra provides a complete visualization and analysis solution for wet laboratory researchers without programming expertise. Figra is freely available and requires no installation beyond the Excel add-in manifest.

## Methods

### Interface design

Figra is designed around a single guiding principle: every step from data loading to figure export should be achievable without writing a single line of code. The Figra control panel is organized into nine thematic tabs, Data & Chart, Converter, Colors & Style, Text & Font, Theme & Layout, Statistics, Analysis, and Help, grouping related controls logically to reduce cognitive load. Users progress naturally from left to right: selecting data and chart type, adjusting appearance, configuring statistics, and exporting.

A key design decision concerns settings consistency. When the user clicks Preview, all current UI settings are captured and stored in memory. Subsequent actions, inserting the figure into Excel, exporting R code, use these stored settings rather than re-reading the UI, guaranteeing that the exported figure and R code always reproduce the exact previewed result.

### Implementation

Figra is built on three core technologies. Office.js (Excel JavaScript API) reads data from the user’s selected cell range and inserts finished figures as images into the worksheet. webR^5^, a port of the R statistical environment to WebAssembly, executes R code entirely within the browser without a local R installation or server connection, ensuring data never leave the user’s machine. All visualizations are generated by custom R functions built on top of ggplot2, with a consistent parameter interface shared across all chart types.

The figure generation pipeline proceeds as follows: the user selects a data range and clicks Load Data; Office.js reads the values and passes them to the JavaScript layer; the data and UI settings are passed to webR as an R function call; R generates an SVG figure using svglite; the SVG is converted to PNG via the canvg JavaScript library; and the PNG is inserted into the Excel worksheet via Office.js.

### Dependencies

Figra relies on ggplot2 as its core visualization engine, executed via webR [6]. Statistical analyses use R’s base stats package (Shapiro-Wilk, t-test, ANOVA, Wilcoxon, Kruskal-Wallis, Friedman, paired t-test, repeated measures ANOVA), supplemented by the car package for Levene’s test, PMCMRplus for Dunn’s test, and multcomp for Dunnett’s test. SVG generation uses svglite. On the JavaScript side, Figra depends on Office.js for Excel integration and canvg for SVG-to-PNG conversion.

## Availability

Figra is freely available at https://h20gg702.github.io/figra-pages/index.html. Users install the add-in by loading the manifest file into Excel, no R installation, no account, and no additional software required. Figra runs on Excel for Windows and Mac.

## Supplementary Tables

**Table S1**. Visual style preset exported by Figra to Excel cells, showing all configurable appearance parameters (font family, font sizes, colors, bar and line dimensions, axis settings, and ggplot2 theme) in a two-column Setting/Value format. Presets can be saved and reloaded within the add-in to maintain consistent figure styling across a manuscript.

**Table S2**. Example of the wide-to-long data format conversion performed by the Converter tab. The original wide-format input (multiple replicate columns per group) is shown alongside the corresponding long-format output (separate Group and Value columns), as exported directly to a new Excel worksheet.

**Table S3**. Statistical analysis output exported by Figra for the bar chart example (Figure 1E). Includes per-group summary statistics, Shapiro-Wilk normality test results, one-way ANOVA result, and Dunnett’s post-hoc comparisons versus the shControl group.

**Table S4**. Statistical analysis output exported by Figra for the grouped line chart example (Figure 1G). Results are reported at each X-axis time point and include per-group summary statistics, Shapiro-Wilk normality test, overall test result (ANOVA or Kruskal-Wallis, selected based on normality assessment), and post-hoc comparisons (Tukey HSD following ANOVA, or Dunn’s test with Bonferroni adjustment following Kruskal-Wallis).

**Table S5**. Grouped IC50 analysis output exported by Figra for the grouped dose-response example (Figure 1J). Includes the estimated IC50 value for each cell line and the full fitted 4PL curve data used to render the figure.

**Table S6**. Linear-quadratic survival analysis output exported by Figra for the grouped LQ example (Figure 1K). Includes α, β, α/β ratio, SF2, D_10_, and D_0_ estimates per group and the full fitted LQ curve data.

